# Time-resolved assessment of single-cell protein secretion by sequencing

**DOI:** 10.1101/2021.12.19.473400

**Authors:** Tongjin Wu, Howard John Womersley, Jiehao Wang, Jonathan Adam Scolnick, Lih Feng Cheow

## Abstract

Secreted proteins play critical roles in cellular communication and functional orchestration. Methods enabling concurrent measurement of cellular protein secretion, phenotypes and transcriptomes are still unavailable. Here, we describe time-resolved assessment of protein secretion from single cells by sequencing (TRAPS-seq). Released proteins are trapped onto cell surface via affinity matrices, and the captured analytes together with phenotypic markers can be probed by oligonucleotide-barcoded antibodies and simultaneously sequenced with transcriptomes. We used TRAPS-seq to interrogate secretion dynamics of pleiotropic cytokines (IFN-γ, IL-2 and TNF-α) of early activated human T lymphocytes, unraveling limited correlation between cytokine secretion and its transcript abundance with regard to timing and strength. We found that early central memory T cells with CD45RA expression (T_CMRA_) are the most effective responders in multiple cytokine secretion, and polyfunctionality involves unique yet dynamic combinations of gene signatures over time. TRAPS-seq presents a useful tool for cellular indexing of secretions, phenotypes, and transcriptomes at single-cell resolution.

## INTRODUCTION

Secreted proteins, such as cytokines, chemokines, cytotoxic molecules, antibodies, and growth factors, make up the secretome and play crucial roles in mediating cell-cell communication and shaping functionality under physiological and pathological conditions^1-5^. Correlation between functional protein release (e.g., cytokines and chemokines) and cellular phenotypes has been extensively studied yet is still inconclusive, especially for immune cells recently undergoing activation/inflammation processes^6,7^. The reasons according for this may include the heterogeneity of cell subpopulations and variable dynamics of protein secretion with regard to timing and strength^8-10^. Moreover, protein secretion is not always consistent with its transcript level that could result from the collective effects of epigenetic^11^ and/or translational/post-translational regulation^12-15^. Therefore, approaches enabling simultaneous measurement of multiple cellular features (e.g., protein secretion, phenotypic traits, and transcriptional profiles) at single-cell resolution, and ultimately being able to achieve time-resolved information of single-cell protein release, will be important not only for the understanding of cell biology but also for the rational design of disease treatment regimens.

Classical methods such as enzyme-linked immunosorbent assay (ELISA) and enzyme-linked immunospot (ELISpot) assess the population-level capacity of protein secretion by measuring extracellularly released analytes without information of source cells producing them^16^. Flow cytometry-assisted intracellular staining assay (ICS) is useful to evaluate single-cell protein expression, which however generally requires trapping of secreted proteins inside the cells prior to fixation/permeabilization at the time point measured, thus unable to represent the true cell secretion and temporal secretion dynamics, also precluding further analysis when live cells matter. Some derivatives of these techniques such as Flow-FISH (combined analysis of cellular protein via ICS and mRNA by probing) and FluoroSpot (fluorescence-based multicolor ELISpot) are still less feasible to determine the temporal dynamics of single-cell protein secretion^17,18^. To meet these gaps, platforms using micro-droplets^19-22^ or microengraved arrays^8,10,23^ are the methods of choice, in which cells are individually compartmentalized and the released proteins are captured onto capture antibodies-coated solid surface (e.g., bead or array slide). In addition, secretion capture assay (SCA), an approach to capturing secreted analytes onto cell surface using affinity matrices such as bispecific antibody that binds to both cell-surface protein and secreted protein, has long been established for the characterization of cytokine-secreting T cells and antibody-expressing B cells^24-26^. While these techniques make it feasible to determine multiplexed and dynamic protein secretion from live single cells with known phenotypes^8,10,21^, they are inevitably hindered by the numbers of secretion targets and surface markers as well that can be concurrently labelled for deep cell characterization due to fluorescence spectral overlap. Moreover, there are limited flexibility and throughput to recover the individual cells of interest for downstream single-cell RNA sequencing (scRNA-seq) even with the possibility using index sorting^27,28^.

scRNA-seq has provided biological insights in a diverse range of research areas and becomes even more powerful with concurrent measurement of multiplexed cell-surface markers using oligonucleotide-barcoded detection antibodies (Ab-Oligo)^29-31^. Inspired by these advances and inadequacies, we describe time-resolved assessment of protein secretion from single cells by sequencing (TRAPS-seq), a method that relies on the time-dependent trapping of secreted proteins on the cell surface and the use of Ab-Oligo barcoding for the quantification of differential protein secretion, enabling cellular indexing of secretion dynamics, cell-surface phenotypic traits, and RNA profiles from sequencing-based readout.

## RESULTS

### Design rationale of the TRAPS-seq

To make protein secretion addressable to its source cell, we took advantage of the design of secretion capture assay (SCA), in which secreted proteins of interest were captured onto the cell surface via affinity matrices such as bispecific antibody that binds to both cell-surface protein and secreted protein (Figure 1A). To enable measurement of protein secretion via sequencing-based readout across thousands of single cells at a time, we adopt oligonucleotide-barcoded detection antibodies (Ab-Oligo) (Figure 1B) that can be easily integrated into current platforms for single-cell multi-omics studies.

**Figure 1.**
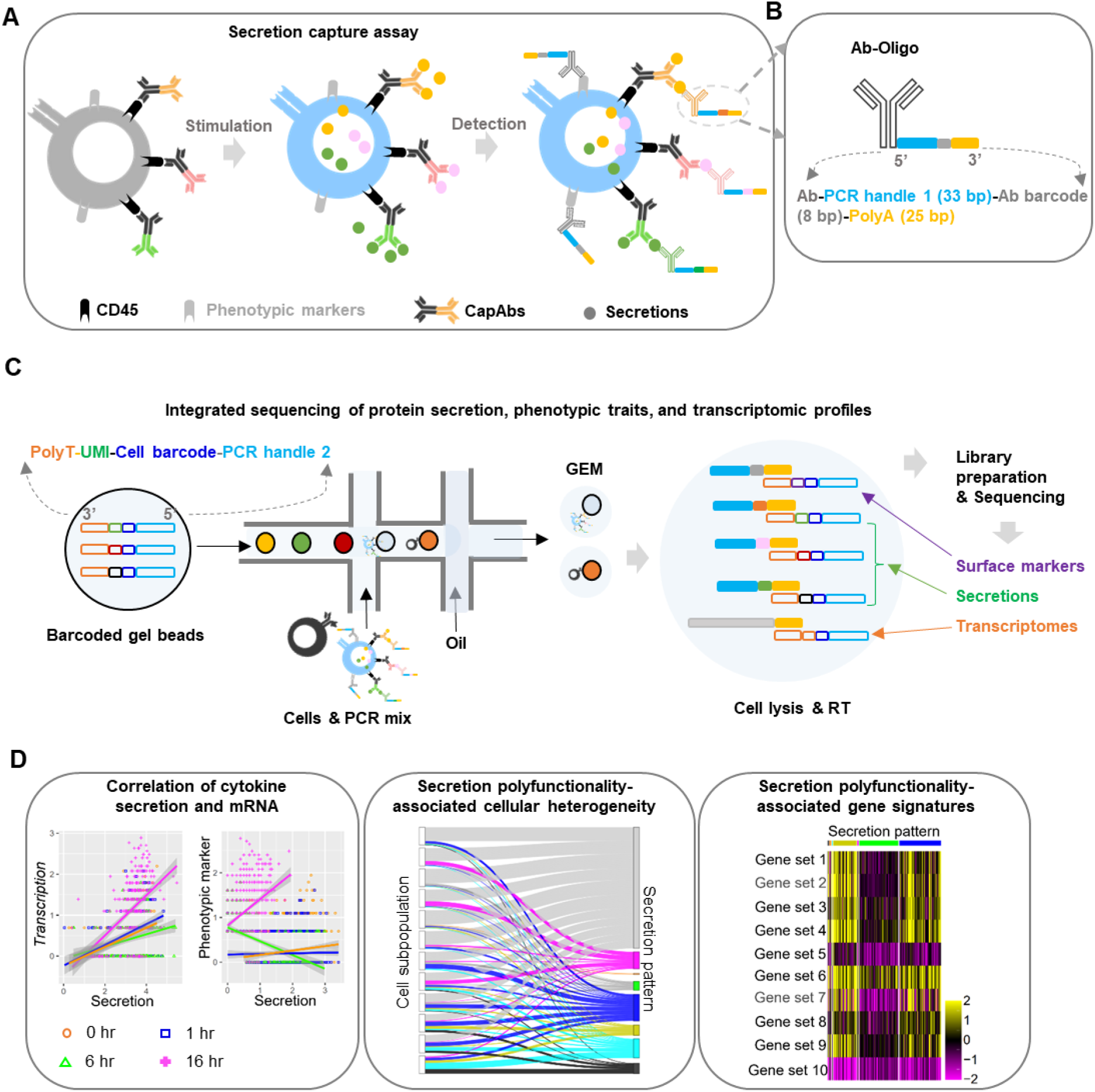
Schematic representation of TRAPS-seq workflow (A-B) Cells were pre-coated with cytokine-capturing matrices (CapAbs) for standard secretion capture assay (SCA) (A) and probed by DNA-barcoded detection antibodies (Ab-Oligo) that may target either the captured cytokines or cell surface markers (B). (C) The stained cells were subjected to droplet-based cell-bead encapsulation and sequencing libraries construction that include mRNA content and barcodes representative of single-cell matched cytokines and phenotypes. (D) Bioinformatics analysis reveals relationship between cytokine secretion, phenotypic traits and transcription state.

The selection of cell-surface anchor sites for cytokine-capturing antibodies (CapAbs) has profound effects on downstream biological studies. CD45, a stable and abundant pan-leukocyte surface glycoprotein is an ideal choice for CapAbs anchoring (Figure S1A). We showed that CD45 staining has negligible effects on T cell expansion capability (Figure S1B). We further demonstrated that the abundance of cell-surface CD45 is not a limiting factor to generation of affinity matrix for multiplexed cytokine-capturing assay (Figure S1C, D). Immediately after the capture of secreted proteins, cells are simultaneously labeled with multiplexed sequencing-compatible Ab-Oligos targeting the surface-bound secreted proteins (Figure S2 and Figure S3) and cell-surface markers (Figure 1C). Subsequent high-throughput single-cell sequencing enables integrated analysis of single cell transcriptome, secretion and phenotypes (Figure 1D).

### TRAPS-seq enables multimodal single-cell characterization

Pleiotropic cytokines (IFN-γ, IL-2 and TNF-α) are key factors orchestrating antigen-elicited immune responses and have been extensively studied^1,2^. However, there is still a lack of our understanding about their secretion dynamics with regard to timing and strength, and also the underling cellular and molecular signatures at a higher dimension.

Herein, we used TRAPS-seq to interrogate the secretion dynamics and functionality of human lymphocytes activated by anti-CD3/CD28 microbeads (Figure 2A), a TCR-dependent stimulation approach used to mimic physiological activation of T cells. Cytokine secretion at time points of 1 hour, 6 hours and 16 hours post cell activation was measured (Figure 2A and Figure S4A). These timepoints were selected to overlap the known expression timing of pleiotropic cytokines (IFN-γ, IL-2 and TNF-α) by T cells^8,32^. The single-cell transcriptomes of unmodified and CapAbs modified lymphocytes was indistinguishable, implying that CapAbs precoating did not perturb cellular transcriptomes (Figure S4B). Activation-induced enhancement or repression of protein secretion was observed for all three cytokines (Figure S4C), suggesting TRAPS-seq was capable of measuring the heterogeneity of protein secretion at the single-cell level. Cells were clustered according to their expression profiles of phenotypic protein markers (Figure S4D) and annotated by using both surface-protein expression and a panel of genes associated with T-cell activation/differentiation (Figure S4E; refer to Figure S6A about background subtraction). Notably, differentiated T-cell subsets and NK-cell subpopulations are clustered more distinctly by surface markers (Figure S5), demonstrating that multimodal analysis of single cells allowed better cell characterization beyond transcriptome alone (Figure S5)^33^.

**Figure 2.**
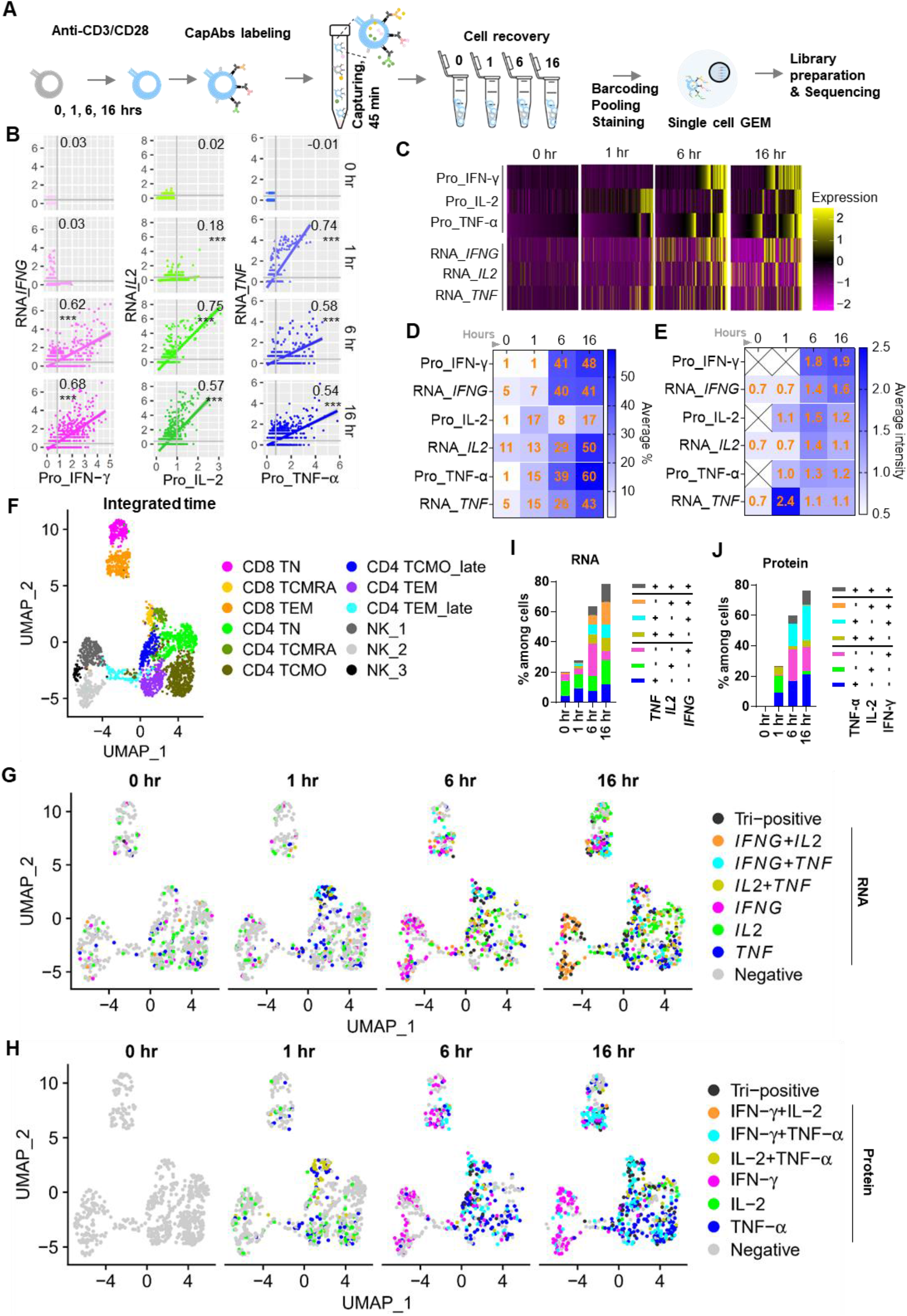
TRAPS-seq enables integrated profiling of single-cell protein secretion, phenotypes and transcriptomes (A) Experimental design of TRAPS-seq using human lymphocytes (B cells-depleted) activated by anti-CD3/CD28 microbeads for indicated time points. (B-C) Dynamic change of cytokine secretion versus transcript level as shown by scatter plots. Shown are log-normalized values and Pearson’s correlation coefficient with significance test. \**P* < .05, \*\**P* < .01, \*\*\**P* < .001 (B) or heatmap at single-cell level over time (C). (D-E) Shown were summary of cell percentile with cytokine releasing or cytokine gene expression and strength, correspondingly. (F-H) Unsupervised cell clustering based on the expression abundance of phenotypic markers (F). The annotated cells were colored by their cytokine gene expression (G) or cytokine release profile (H). Shown histogram plots were summary of cytokine expression polyfunctionality corresponding to G (I) and H (J).

We found that there was limited correlation between cytokine secretion and gene expression (refer to Figure S6B, C about background subtraction) with regard to timing and strength (Figure 2B-E and Figure S7), and this inconsistency was also true for the surface proteins detected (Figure S7B)^29,34^. TNF-α and IL-2 secretion occurred earlier than IFN-γ following cell activation (Figure 2C, D). For long-term activation (6-16 hours), the percentage of IL-2-positive cells remained relatively stable while expression level of *IL2* mRNA continuously increased. On the other hand, both the proportion of cells and gene expression levels of cells secreting TNF-α or IFN-γ increased over time (Figure 2D). TNF-α or IL-2 secretion was higher than IFN-γ in response to initial stimulation (1 hour) but diminished at late activation stages (6-16 hours), while IFN-γ secretion per cell gradually increased (Figure 2E). This is in line with the gradual functionality acquisition of T/NK cells upon activation and differentiation^9,35,36^. These data suggest that time-resolved measurement of both protein secretion and transcription state is more insightful in our understanding of cellular function acquisition.

A deep mapping of cytokine mRNA and protein onto cell subtypes revealed high diversity of cells in cytokine production (Figure 2F-H). This heterogeneity was cell phenotypes and/or activation state-related, resulting in discordance in both the timing and multiplicity between cytokine gene generation and protein release (Figure 2I, J). Intriguingly, NK cells had a sparse secretion of IL-2 post initial activation and shifted to IFN-γ during late stage, the gene expression, however, possessed a distinct pattern from *IFNG* to *IFNG*^+^ *IL2*^*+*^ (Figure 2G, H). The presence of *IL2* mRNA in NK cells is controversial but could be culture-system varied^37,38^.

### TRAPS-seq identifies cell subpopulations with heterogeneous secretion potency

The presence of polyfunctional cytokine-producing T cells has been associated with protective immunity^39,40^ and favorable therapeutic outcomes^41^. A comprehensive understanding of the occurrence and maintenance of these heterogeneous cells at both cellular and molecular levels would be invaluable.

Thus, we further dissected the secretion potency of heterogeneous cell subpopulations, especially T lymphocytes. We found that pre-existing central memory T cells with CD45RA expression (T_CMRA_) (Figure S4E)^42,43^ regardless of CD4^+^ T or CD8^+^ T cells had the rapidest cytokine secretion, primarily TNF-α^+^ IL-2^+^ and TNF-α^+^ within one hour post activation, while those more-differentiated counterparts like central memory without CD45RA expression (T_CMO_) and effector memory (T_EM_ and T_EM_late_) tended to secret mono-cytokine such as TNF-α^+^ or IL-2 during initial stimulation (Figure 3A). CD8^+^ T_CMRA_ continued to release multiple cytokines, specifically IFN-γ-involved triple or double positive cytokines, whereas amongst CD4^+^ T cells, CD4^+^ T_CMRA_ and CD4^+^ T_CMO_late_ were the multi-cytokine releasing cells (Figure 3B, C). This discrepancy could result from activation-coupled subtler phenotypic change from CD4^+^ T_CMRA_ to CD4^+^ T_CMO_/CD4^+^ T_CMO_late_ during stimulation^39^. On the other hand, CD8^+^ T_EM_ and NK cells had mainly contributed to the mono-IFN-γ expression per cell during late activation stage (Figure 3B, C). Naïve T cells (T_N_) instead were shown to be more inert to CD3/CD28-engaged activation but endowed with the highest capacity in secreting various combination of cytokines for a long term stimulation (Figure 3C), consistent with their developmental diversity post exposed to exogenous antigens^44^.

**Figure 3.**
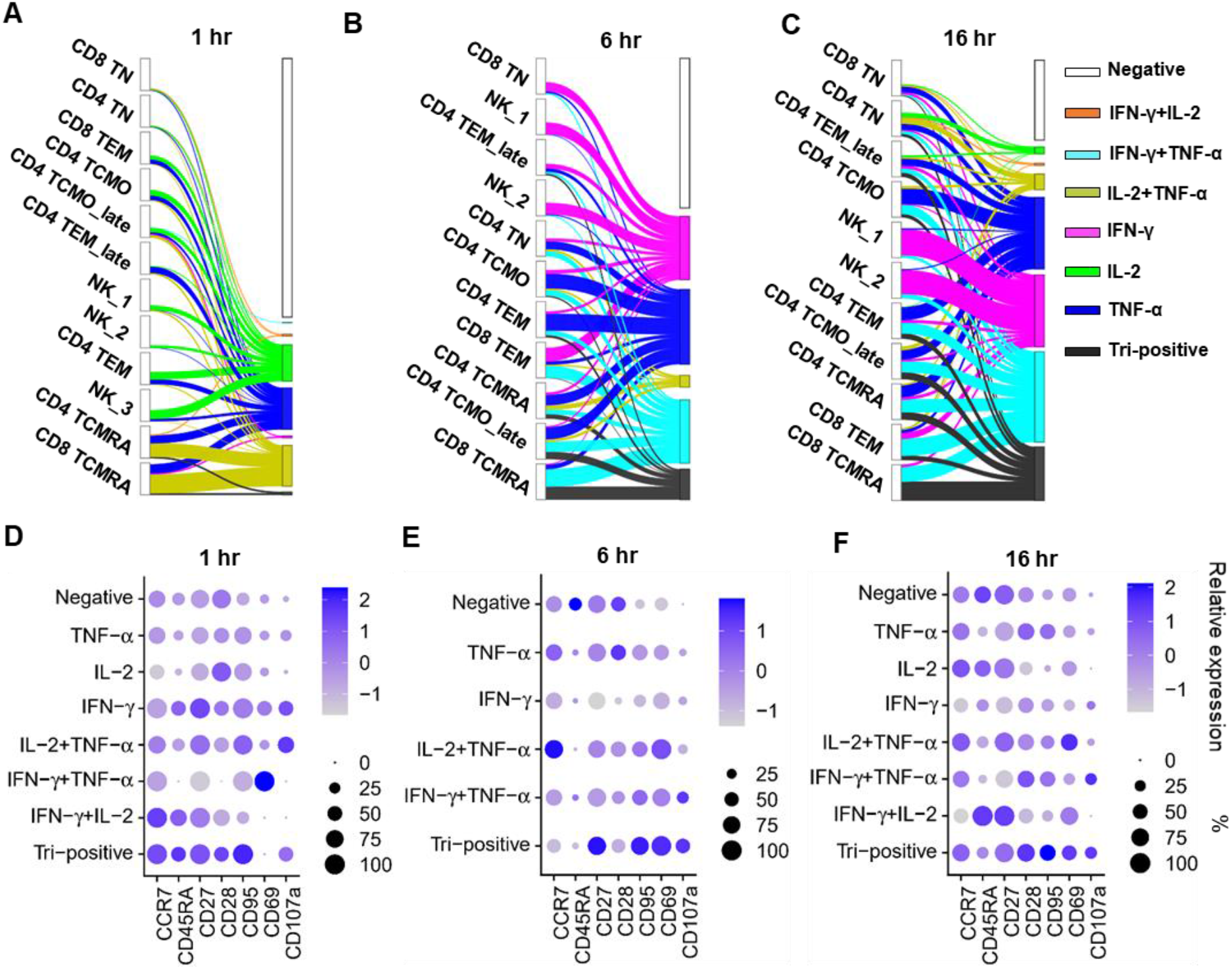
TRAPS-seq identifies T_CMRA_ as the most effective responders in multiple cytokine secretion (A-C) Sankey diagrams depict how cell composition/activation state change over time may affect their cytokine secretion profiles. Cell numbers for each cell type were normalized. (D-F) Expression of a panel of phenotypic markers in T cells with indicated cytokine release profile.

While there was limited correlation between individual cytokine secretion and surface marker expression (Figure S8), combinatory analysis of surface phenotypic/activating markers could partially identify cells of distinct secretion potency (Figure 3D-F). T cells with triple cytokine release at 1 hour post stimulation had stem-cell memory (T_SCM_)-like phenotypic markers (e.g., CCR7, CD45RA, CD27, CD28 and CD95)^43^ (Figure 3D). Late activating T cells with triple cytokine release additionally had high abundance of the activation marker CD69 and degranulation molecule CD107a (Figure 3E, F). Cells capable of secreting IL-2^+^ TNF-α^+^ were characterized by high CD107a and to a less extent CD95 in their resting/early state, and both of which were diminished along with the elevation of CD69 for a long-term activation, however, cells producing IFN-γ^+^ TNF-α^+^ were characteristic of the opposite dynamics (Figure 3D-F). Therefore, cytokine secretion potency of CD3/CD28-engaged T cells was cell subtype associated and affected by their dynamic change of phenotypic/activating markers, but less correlated to change of a singular marker.

### TRAPS-seq characterizes transcriptional signatures of cytokine secretion polyfunctionality

Single-cell RNA sequencing-derived transcriptome profile has been widely used for the study of polyfunctional T cells, however, being with a lack of real cytokine secretion. We next attempted to characterize the transcriptional signatures that may be correlated with cytokine secretion potency.

Notably, despite limited correlation between cytokine secretion and mRNA abundance (Figure 2B, C and Figure S7), *IFNG, IL2*, and *TNF* were still among the top genes that differentiated cells with or without that cytokine secretion, correspondingly (Figure 4A, B). *FOS* (AP-1 transcription factor subunit), *EGR1* (early growth response protein 1) and *CD69* (early activation marker) were enriched in cells secreting IL-2^+^ TNF-α^+^ and to a lesser extent TNF-α^+^ only, but not IL-2^+^ (Figure 4A), consistent with the finding that IL-2^+^ TNF-α^+^ /TNF-α^+^ was among the first wave of stably released cytokines during T-cell early activation (Figure 3A). Cells with IFN-γ-involved cytokine release were distinguishable from those without IFN-γ secretion, manifested by an upregulation of *IFNG* but also genes encoding chemokine ligands (e.g., *CCL4, CCL3, CCL4L2, XCL2*, and *CCL5*) and cytotoxic molecules (e.g., *GZMB* and *NKG7*) (Figure 4A, B), indicating their strong effector function. Cells with triple cytokine secretion had a general downregulation of chemotactic and cytotoxic mRNAs (e.g., *CCL4, CCL3, CCL4L2 NKG7, GZMB* and *GNLY*) compared to IFN-γ^+^ or IFN-γ^+^ TNF-α^+^ cells (Figure 4B). However, triple cytokine positive cells shared some unique features with those secreting early cytokines IL-2^+^ TNF-α^+^/TNF-α^+^, for example, an upregulation of genes relating to immune and inflammatory responses such as *CSF2* (granulocyte-macrophage colony-stimulating factor), *IL21* (Interleukin 21), *IL1R1* (interleukin 1 receptor, type I) and *IRF8* (interferon regulatory factor 8), and meanwhile, genes favoring cell division like *IER3* (immediate early response 3, protecting cells from Fas- or TNF-α-induced apoptosis), *CCND3* (G1/S-specific cyclin-D3) and *EZH2* (enhancer of zeste homolog 2, critical for epigenetic modification) (Figure 4A, B), suggesting an unique state of this triple cytokine-secreting cell subpopulation which was endowed with not only immediate/regulatory functions but also superior commitment to ensuing proliferation/differentiation^39,45^.

**Figure 4.**
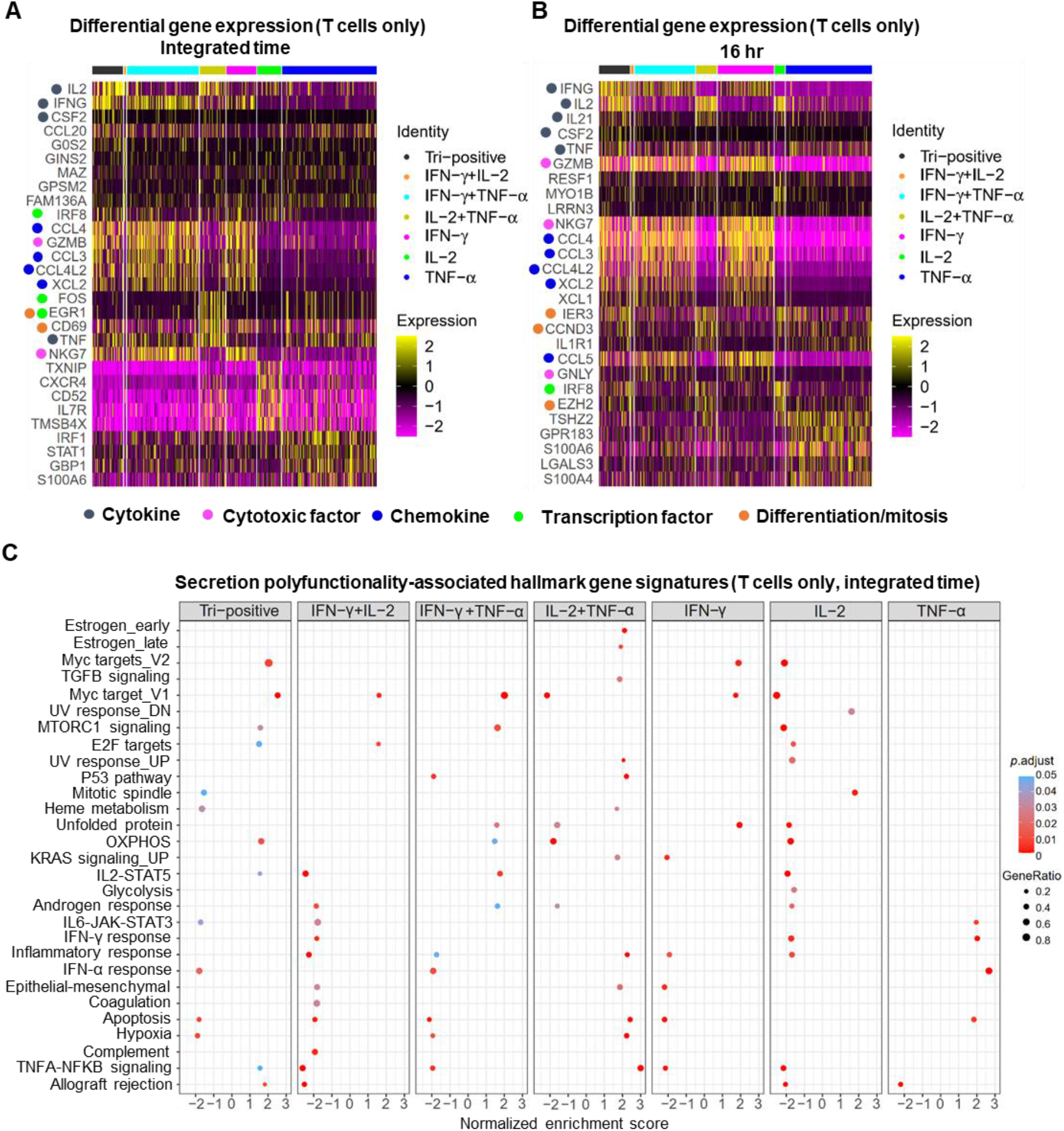
Cytokine-secreting polyfunctionality-associated differential gene-set signatures (A-B) Differential gene expression of T cells with indicated cytokine release. Shown were top five differentially expressed genes for cells from integrated time points (A) and time point of 16 hours (B) post activation. (C) Identified hallmark gene set signatures distinguishing T cells of varied cytokine-secreting potency.

To depict the whole landscape of T-cell cytokine secretion and transcriptome profile, we calculated gene set-based scores at single-cell level using the curated hallmark gene sets from the Molecular Signatures Database (MSigDB). We found that IL-2 and TNF-α signaling were highly engaged in IL-2/TNF-α-involved polyfunctional cells but not those secreting IL-2 or TNF-α only, implying that effective acquisition of multi-cytokine secretion could involve on both IL-2- and TNF-α-mediated paracrine/autocrine effects (Figure S9A-C). On the other hand, IFN-γ signaling was more engaged in cells secreting TNF-α only, suggesting potential regulation on TNF-α secretion via IFN-γ-mediated paracrine effects (Figure S9A, D). Moreover, cells with triple cytokine secretion had a generally higher enrichment of genes involved in oncogenic transcription factor Myc-regulated targets and oxidative phosphorylation (OXPHOS), but a downregulation of genes relating to IFN-α response, apoptosis and hypoxia (Figure 4C and Figure S9E), a favorable cellular state for optimal cytokine secretion and survival. Of note, despite that the expression timing and strength of genes representative of known signatures could be dynamically varied over time, it was still feasible to characterize cells of discernable secretion potency by a combination of certain unique gene set-based signatures for each individual time points (Firue S9F-H).

## DISCUSSION

In this study, we describe for the first time a sequencing-based approach to measuring dynamic protein secretion at single-cell level (TRAPS-seq). We have also demonstrated how TRAPS-seq can be simultaneously integrated with sequencing of transcriptomes and phenotypic proteins to dissolve the cellular and molecular heterogeneity of differential cytokine secretion.

We found that there was limited correlation between cytokine protein secretion and mRNA expression for all cytokines detected (IFN-γ, IL-2 and TNF-α) in CD3/CD28-engaged T cells. Combined analysis of single-cell intracellular cytokine abundance and mRNA level (Flow-FISH), however, showed that the expression dynamics of intracellular cytokine protein was generally consistent with its mRNA^18^. This discrepancy could mainly result from the measurement of cytokine protein expression by TRAPS-seq is a snapshot compared to progressive accumulation (with the protein transporter machinery inhibited) by Flow-FISH at the time point measured. The disconnect of cytokine secretion and transcription has profound effects on protective and dysfunctional immune responses (e.g., autoimmune diseases and inhibited antitumor activity)^14,15,46^. TRAPS-seq, when combined with single-cell transcriptome measurement, would provide us with more comprehensive information to advance our understanding of cytokine-involved physiological and pathological processes.

T_CMRA_ cells are phenotypically resembling T_SCM_ (or T_SCM_-like) ^43,47^, characterized by positive expression of canonical naïve/memory-specific markers CCR7 and CD45RA (but lower than T_N_), and high expression of Fas receptor CD95 and cell persistence-related costimulatory molecule CD27. T_CMRA_ has also a highly presented lysosomal-associated membrane protein-1 (LAMP-1/CD107a), suggesting a potential state of T_CMRA_ (especially CD8^+^ T_CMRA_) ready for immediate cytotoxic function. We have also demonstrated that T_CMRA_ was the most effective responder in secreting ample amount of IL-2 and TNF-α or TNF-α only soon post CD3/CD28 engagement and towards the secretion of triple cytokines after a longer activation course. These IL-2^+^ TNF-α^+^ or IL-2^+^ TNF-α^+^ IFN-γ^+^ had a dramatic upregulation of genes involved in *IL2-STAT* and *TNF-NFKB* signaling pathways, and genes favoring cell survival and proliferation/differentiation (e.g., *CSF2, IER3, CCND3, EZH2*, and Myc-regulated genes). These data have extended our previous finding that the generation of T_CMRA_ during the production of therapeutic T cells was well-correlated with the proportion of T_N_ in cellular starting material^42^, and cell products containing more T_CMRA_ were manifested by younger phenotypes and superior functions that are recognized to be good for therapeutic outcomes. Thus, combined analysis of secretion-included single-cell multi-omics would see their versatility in the characterization of secretion profiles and dynamics that could be associated with protective immunity or harmful inflammation, facilitating the rational design of therapeutic regimens^7,41,48^.

Finally, despite its usefulness of TRAPS-seq, a limitation of this strategy could be cytokine capture multiplexing. To circumvent this, additional common anchoring sites of CapAbs could be further explored beyond CD45 (used in this study), for example, via chemical modification of cell surface to increase artificial binding sites^49^. Meanwhile, DNA barcode design with enhanced signal-to-noise ratio would also be helpful^50^. Undoubtedly, TRAPS-seq has filled up a gap in the area of single-cell multi-omics studies^51^, making cytokine secretion indexable to multi-modalities of its source cell, and would find its versatility in near future.

## SUPPLEMENTARY INFORMATION

Supplementary information is available for this manuscript.

## ACKNOWLEDGEMENTS

This work was supported by funding from Singapore Ministry of Education Academic Research Fund Tier 2 (MOE-000063) and Institute for Health Innovation and Technology (iHealthtech), NUS. The authors acknowledge technical support from the Flow Cytometry Laboratory (NUS Medical Sciences Cluster) for cell sorting.

## AUTHOR CONTRIBUTIONS

T.W and L.F.C conceived and designed the study. T.W performed the experiments and data analysis with input from L.F.C and J.A.S in data mining and assistances from H.J.W and J.W in sequencing techniques. T.W and L.F.C wrote the manuscript with feedback from J.A.S. All authors approved the submission.

## METHODS

### Cells and reagents

Apheresis blood samples from healthy donors were gifts from the Health Sciences Authority (HSA), Singapore, with approval from Institutional Review Board and informed consent from donors (NUS-IRB no. H-18-038E). Ficoll Paque Plus (GE Healthcare, 17-1440-02) was used for peripheral blood mononuclear cells (PBMC) separation. RPMI1640 medium (ThermoFisher, A1049101) with 10% fetal bovine serum (FBS) (ThermoFisher, 10270106) was used in cell culture. Antibodies and Ab-Oligo used in this work were listed in Table S1 and DNA oligos for antibody modification and q-PCR/sequencing were synthesized by IDT (Integrated DNA Technologies) and listed in Table S2. Cytofix/Cytoperm reagent kit (BD Biosciences, 554714) was used for intracellular cytokine staining assay. For immunostaining, Fc-blocking reagent (Biolegend, 422301) was used to block Fc receptor-introduced non-specific staining. Phosphate buffered saline (PBS) with 2% FBS was used for staining buffer and 0.5% FBS for washing buffer, except where specified.

### Ab-Oligo conjugates generation and validation

Ab-Streptavidin conjugates were generated according to the instruction of Lightning-Link Streptavidin Antibody Labeling Kit (Novus Biologicals, 708-0030). Synthesized Oligo-biotin was conjugated with Ab-Streptavidin at a saturated biotin-to-streptavidin molecular ratio (8:1) at room temperature for 90 min. Excess free oligo-biotin was removed by streptavidin-coated microbeads (Invitrogen, 65001) and validated by polyacrylamide gel electrophoresis (PAGE). The produced Ab-Oligo complex was stored in PBS buffer containing 0.5% FBS, 1 mM EDTA, and 0.09% NaN_3_.

Unstimulated PBMCs were stained with cytokine-capturing CapAb and incubated in cytokine-containing culture supernatant at 4°C for 20 min. For control group, cells were incubated in staining buffer instead. Post thorough washing the cells were further stained by indicated Ab-Oligo (0.3 – 0.5 µg/test) separately at 4°C for 30 min and probed by complementary FAM-polyT primer (2 µM) which binds to the polyA tail of antibody barcodes at room temperature for 20 min. Meanwhile, the competitively blocking function of modified Ab-Oligo conjugates was validated by their capability to reduce the binding of fluorescent antibodies (Ab-Fluo) of the same antibody clones.

### Cytokine secretion capture assay (SCA)

Cytokine SCA was carried out according to the standardized protocol with the kits (Miltenyi Biotec) as listed in Table. S1, but with slight modification. Briefly, CD19^+^ cells-depleted lymphocytes were sorted in Moflo Astrios cell sorted (Beckman Coulter) and kept in incubator overnight before stimulation. Cells were activated by anti-CD3/CD28 Dynabeads (Gibco, 11161D) at 2:1 bead-to-cell ratio for indicated time periods in 96-well U-bottom plate. Post beads removal, cells were stained with saturated cytokine-capturing antibodies for 15 min at 4°C and subjected to secretion capture at 37°C for 45 min with gentle rotation. The captured cytokines can be detected by fluorescent antibodies for flow cytometry analysis or by Ab-Oligo for downstream sequencing experiments.

### Sample staining with pooled Ab-Oligo

Ab-Oligo for cell surface markers (1 µg/test) and captured cytokines (0.3-0.5 µg/test) were pooled and filtered by Amicon Ultra-0.5 centrifugal column (MWCO 100 kDa) immediately before use. The filtered Ab-Oligo mixture was re-suspended in 50 µl PBS with 2% heat-inactivated FBS. Cells for each time points (0, 1, 6, 16 hours) were individually labelled with β2M-targeting hashtags sand Fc-blocking agents, while indicated, control cells without CapAb modification were labelled with CD45-targeting hashtags instead. These five group of cells were pooled together after washing and stained with Ab-Oligo cocktail for 45 min at 4°C. Cells were washed with cold PBS containing 2% FBS for 4 times and re-suspended in PBS before run on the Chromium Controller (10x Genomics).

### Sequencing libraries preparation for TRAPS-seq

The 10x Genomics single cell 3’ reagent kits (v3.1 Chemistry) was used to generate single-cell barcoded initial sequencing libraries following reverse transcription that include complementary DNA (cDNA) of mRNA and protein libraries representative of cytokines, surface markers and sample-indicating hashtags. Post cleanup using silane beads, 35 µl of the eluted DNA sample was pre-amplified for the 1^st^-round following the standard protocol with the addition of two additive primers for surface protein and hashtag (listed in Table S2) and cycled as follows: as follows: 98°C 3 min, 12 cycles of: 98°C 15 s, 63°C 20 s, and 72°C 1 min; Then an extension step of 1 min at 72°C. The 1^st^-round PCR product was cleaned up and followed by a double size selection step using SPRIselect reagent (Beckman Coulter) to separate protein library from cDNA library.

The amplified cDNA product was subjected to final gene expression library construction according to the instructions with the kit. Part of the protein DNA product (5 µl) was used for the preparation of sequencing libraries corresponding to surface marker and cytokine while additional part (5 µl) was used for the construction of sequencing library of sample hashtags using KAPA HiFi Master Mix (Roche). The 2^nd^-round P7/read 2-incorporated PCR primers used for protein library preparation were listed in Table S2 with the P5/read 1-incorporated primer the same to that used in cDNA library. The size range and concentration of final library constructions was verified by Agilent 2100 Bioanalyzer.

### Single-cell RNA and protein data processing

The libraries for cDNA and sample hashtag were pooled and sequenced in Illumina HiSeq platform by Novogene. The protein libraries for surface protein and cytokine were sequenced in MiSeq platform using MiSeq Reagent Kit v3 and customized sequencing primers (listed in Table S2). Raw cDNA FASTQ data was processed using Cell Ranger v4.0 (10x Genomics) with default parameters. A total of 4,899 cell barcodes with 90,834 reads/cell were reported. These cDNA library-derived cell barcodes were used to match the unique molecular identifier (UMI) counts of sample hashtags, surface protein and cytokine using CITE-seq-Count with a Hamming distance of 1 for both cell barcodes calling and UMI counting^52^. Pre-processed count matrices were imported into Seurat package (v4.0)^33^, and dead cells were removed by filtering out cells expressing > 5% mitochondrial transcript counts. Post cell demultiplexing, a total of 4,010 singlets were remained for downstream analysis. For the analysis of cytokine secretion, where indicated, sparse dots above 99 quantiles of the background-counts distribution at “0 hr” (Figure S6C) and apparently discrete dots within data of other time points (Figure S7A) were removed.

### Gene set enrichment analysis (GSEA)

Differential gene expression by Wilcoxon Rank Sum test between cells of distinct secretion pattern were performed in Seurat (v4.0) using sctransform-normalized gene expression counts^33,53^, and generated gene lists were imported for GSEA using R programs embedded in the clusterProfiler through the use of curated hallmark gene sets from the Molecular Signatures Database (MSigDB, v7.4)^54,55^. Normalized enrichment scores (NES) and adjusted *P*-value (< 0.05) were ranked for presentation.

### Single-cell gene set-based signatures’ calculation

Single-Cell Signature Explorer was used to score gene set-based signature for each individual cells^56^. Briefly, sctransform-normalized gene expression matrix corresponding to each individual time points was imported into the Single-Cell Signature Scorer to compute signature score per cell over a list of hallmark gene sets from MSigDB (v7.4). Dimensionality reduction plot corresponding to each individual time point was produced by UMAP based on the cytokine secretion measured. The calculated signature scores for individual cells were visualized on the UMAP plot via Single-Cell Signature Viewer. Differentially expressed gene-set signatures (average logFC > 0.25; adjusted *P* < 0.05; 20% of cells within the cluster expressing the target gene) across cell groups with different secretion patterns were performed by Seurat (v4.0) using a Wilcoxon Rank Sum test and shown by heatmap plots^33^.

### Pearson correlation coefficient test

To make it quantifiable and comparable between protein and mRNA expression or protein and protein expression, the varied background signaling of surface marker and cytokine staining was subtracted (refers to Figure S6A, C). For the Pearson’s correlation analysis, where indicated, either all cells or cells that had at least one positive expression of protein or mRNA were included. Paired Pearson’s correlation between protein and mRNA or cytokine release and surface protein marker was performed using correlation package^57^.

### Data analysis

Flow cytometry data were acquired in CytoFLEX (Beckman Coulter) and analyzed using FlowJo software (Tree Star). Histogram or bar graphs, except indicated, were generated in GraphPad Prism software.

## DATA AND CODE AVAILABILITY

The data that support the findings of this study can be made available by the corresponding author.

