## Supplemental File for "Time-resolved assessment of single-cell protein secretion by sequencing"

### Supplementary methods

#### CD45 abundance and stability on lymphocytes

PBMC were recovered overnight before stimulated with or without 50 ng/ml phorbol 12-myristate 13-acetate (PMA) plus 1 μg/ml ionomycin for 12 hours. The cells were collected and pre-stained with saturated concentration of PE-labelled anti-CD45 at 4°C for 20 min. After thorough washing, cells were split into 96-well tissue culture plate at 0.1 million cells/well and cultured for additional hours as indicated in the presence of PMA and ionomycin. At indicated time points, cells were collected and stained with Live/Dead viability dye and fluorescent anti-CD3 for T cell identification. The data was acquired in flow cytometer CytoFLEX (Beckman Coulter).

#### Cell proliferation assay

PBMCs were pre-stained with proliferation-tracing dye (Invitrogen, C34564) following the standardized protocols with the kit. After washing, cells were split into two groups and incubated with or without anti-CD45 (clone HI30, 10 μg/ml) at 4°C for 20 min. Cells were then cultured in 96-well plate with the ratio of anti-CD3/CD28 Dynabeads (Gibco, 11161D) to cells at 3:1 in condition of 5 ng/ml human recombinant IL-2 (ThermoFisher, PHC0021). At day 5 post activation, cell proliferation was analyzed by flow cytometry.

**Preparation of cytokine-containing culture supernatant**

PBMC were seeded at $3x10^6$ cells/well in 6-well plate and stimulated with 50 ng/ml PMA plus 1 µg/ml ionomycin for 10 hours. The culture supernatant was pooled together and centrifuged at 700 g for 10 min to remove cell debris. The supernatant was split into small aliquots and kept at - 80$^o$C.

**Antibody barcodes quantification by quantitative PCR (q-PCR)**

Quantitative PCR was used to correlate the amount of Ab-Oligo to secretion level. To artificially generate cell populations bearing gradient IFN-γ concentration, unstimulated cells were pre-coated with IFN-γ CapAb and incubated in IFN-γ-containing supernatant for different time period. After washing, the cells were stained with 1 µg oligo-barcoded anti-IFN-γ and followed by probing with FAM-polyT primer to indicate the abundance of cell-bound anti-IFN-γ-Oligo. For each condition, a total of 8 single cells were sorted into PCR tube by FAM intensity in Moflo Astrios cell sorter (Beckman Coulter). The sorted cells were directly subject to antibody barcodes quantification by q-PCR. In real secretion capture assay (SCA), PBMCs were pre-stimulated by PMA and ionomycin for 6 hours before subjected to SCA. Finally, cells were co-stained with fluorescent antibodies (Ab-Fluo) and Ab-Oligo of the same antibody clone for each target, the latter of which was further probed by FAM-polyT primer. For each group, three populations of cells and 8 single cells each were sorted according to Ab-Fluo intensity. Q-PCR was applied to correlate the fluorescence intensity with barcode abundance.

**Q-PCR**

Cells were directly sorted into PCR tube containing 14 µl 1 x SYBR Green PCR master mix (Bio-Rad, 1725271) and primer set targeting the PCR handles located on the antibody barcode and FAM-polyT. In the first round PCR, a simple extension step at 72$^{o}$C for 15 s was done to make full complementary dsDNA template formed by FAM-polyT and Oligo. During the second round PCR, normal PCR cycles (40 cycles) was run to amplify the templates in the Agilent AriaMx Real-Time PCR System. For preparation of DNA standards, known concentration of oligo barcode and FAM-polyT were mixed at 1:1 and annealed at 56$^{o}$C before further extended at 72$^{o}$C for 5 min to form dsDNA. The dsDNA standards were then serially diluted, and unstained cell lysate equal to single cell was spiked into each reaction as standard control.

91 ## Supplementary Figures

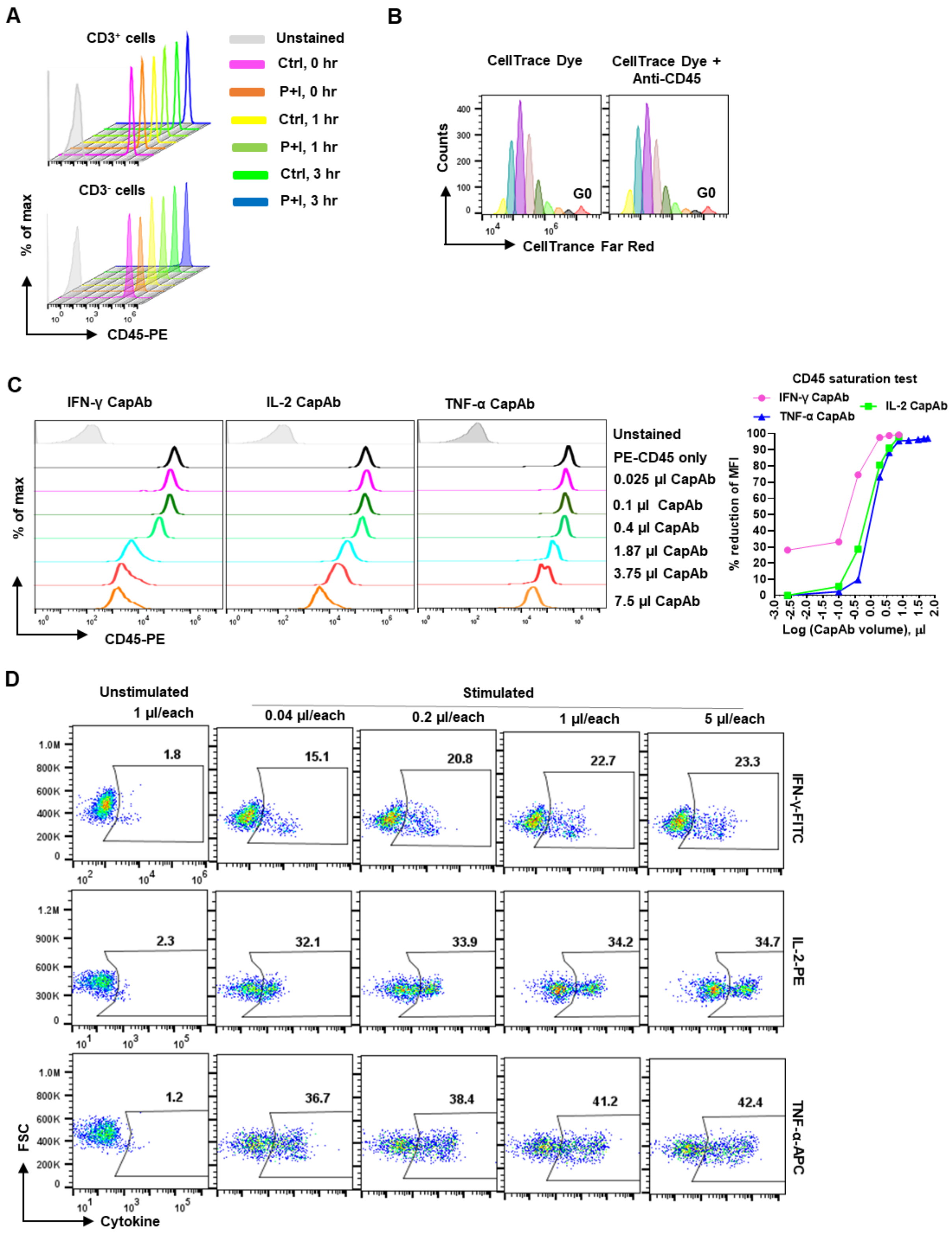


92

**Figure S1. Feasibility test of CD45-targeted secretion capture assay**

(A) Pre-treated ("P+I") or non-treated ("Ctrl") PBMC using PMA and ionomycin for 12 hours were stained with anti-CD45-PE and further cultured in the presence of PMA and ionomycin. PE signaling on cell membrane was detected to indicate the abundance and stability of CD45 molecule during activation. (B) Perturbation on cell proliferation capability post CD45 engagement by anti-CD45 monoclonal antibody. (C) Saturation test of cytokine-capturing antibodies (CapAb) by competitive binding assay using anti-CD45-PE and CD45-targeting bi-specific CapAbs. (D) Multiplex secretion capture assay using serial dilution of CapAbs.

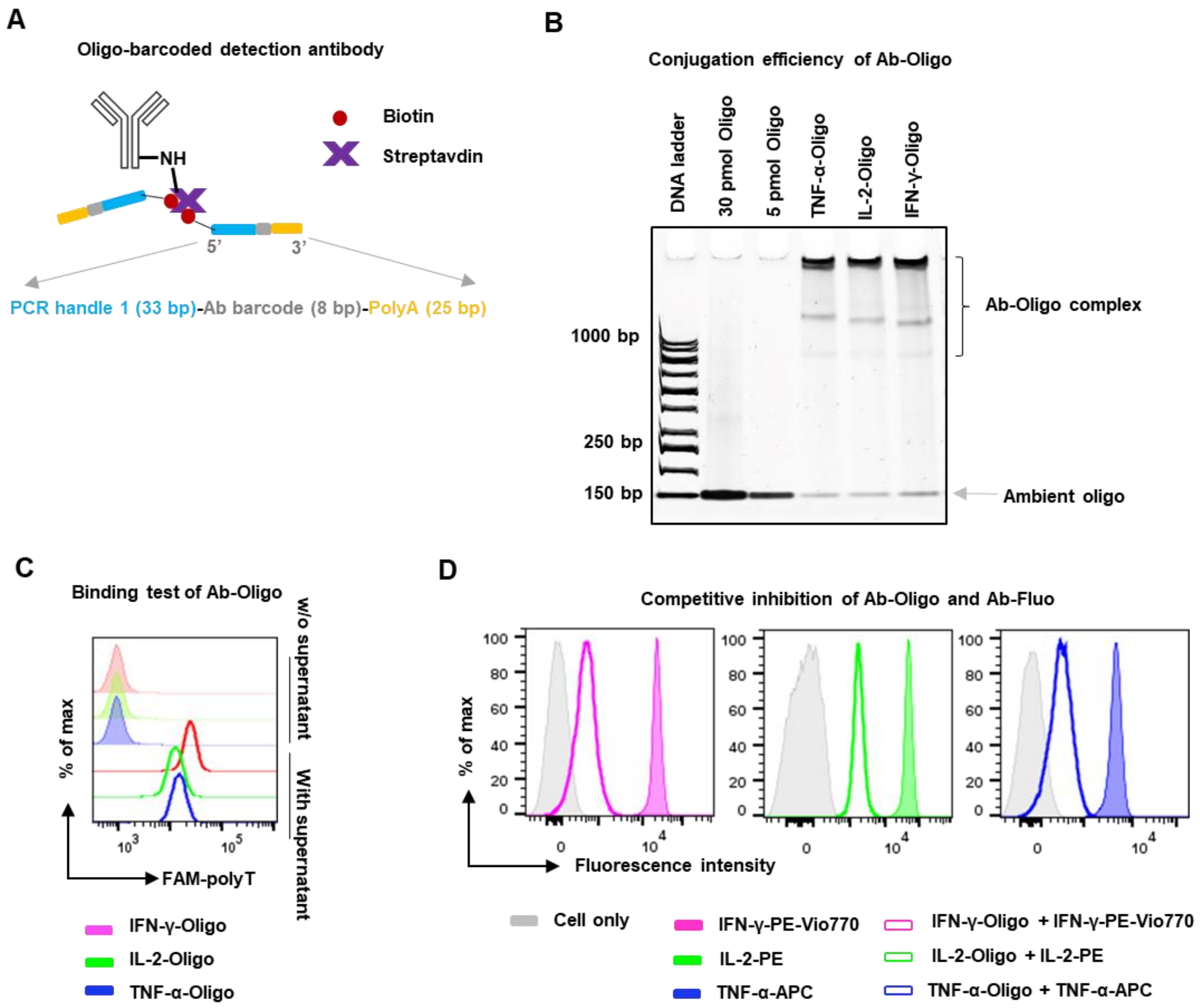


**Figure S2. Generation of oligo-barcoded cytokine detection antibodies**

(A-B) Antibodies were covalently coupled with streptavidin before conjugated with oligo barcodes (A). Shown PAGE suggests successful conjugation of Ab-Oligo and maximal clean-up of ambient oligo (B). (C-D) Functional maintenance of modified antibodies were evaluated. Unstimulated CapAbs-coated PBMCs were pre-incubated in cytokine-containing culture supernatant (SU) before further stained with Ab-Oligo and detected by complementary FAM-polyT primer (C). Also, competitively blocking function of modified antibodies was validated by their capability to inhibit the binding of fluorescent antibodies of the same antibody clones (D).

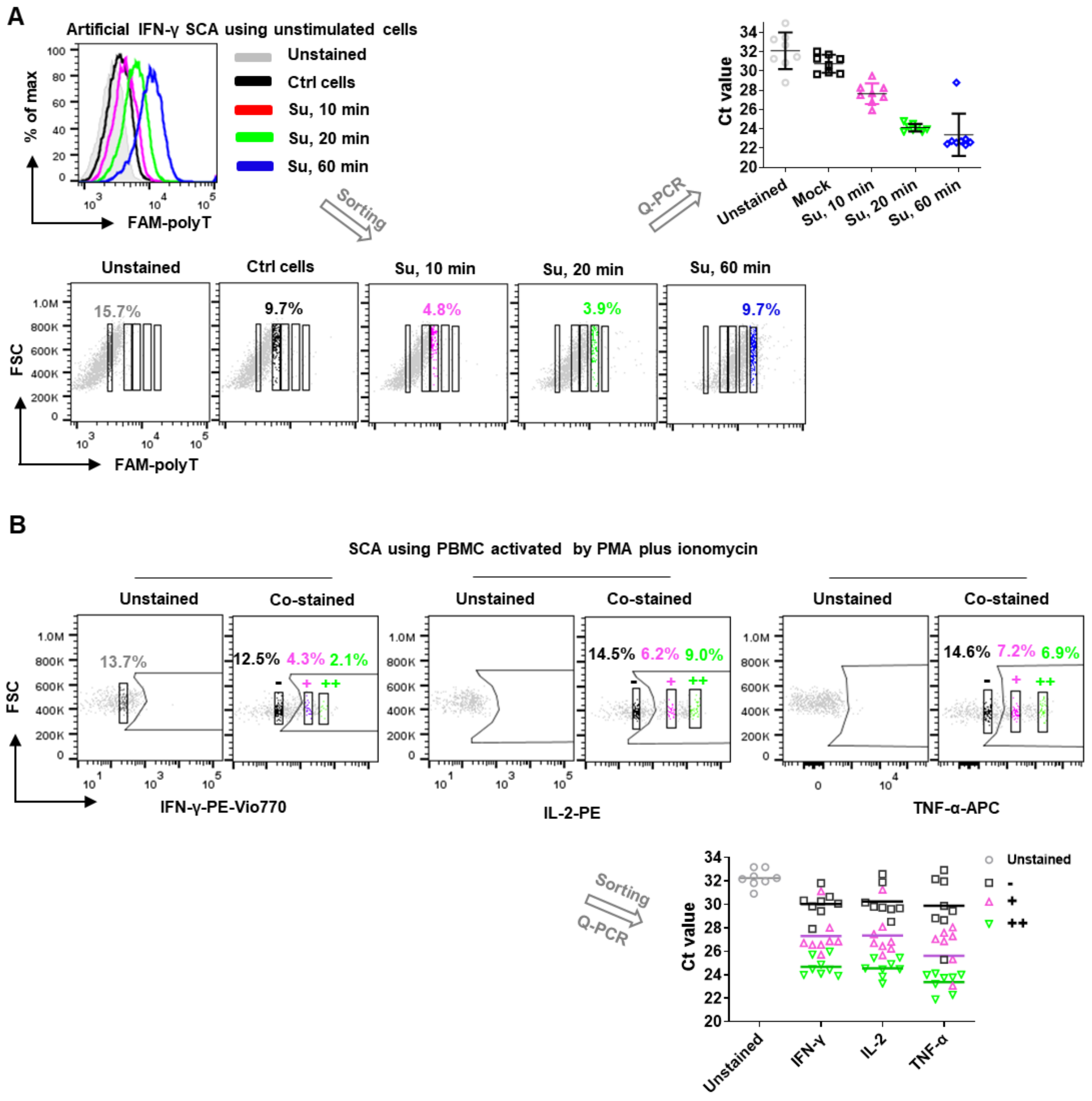


**Figure S3. Use of oligo-conjugated antibodies as molecular counter of cytokine secretion**

(A) Cells were pre-coated with IFN-γ-specific CapAbs and incubated in IFN-γ-containing supernatant (Su) for varied time period to artificially generate cell populations bearing gradient IFN-γ concentration. The cells were stained with FAM-polyT primer to indicate the existence of anti-IFN-γ Ab-Oligo. For each population, a total of 8 single cells were sorted by FAM intensity and semi-quantified by q-PCR. (B) In real SCA, PBMCs were pre-stimulated by PMA for 6 hr before subjected to SCA. Cytokine capturing were detected by co-staining of fluorescent antibodies (Ab-Fluo) and Ab-Oligo. For each group, three populations of cells and 8 single cells each were sorted based on the fluorescence intensity of the Ab-Fluo. Q-PCR was used to correlate the fluorescence intensity to barcode counts.

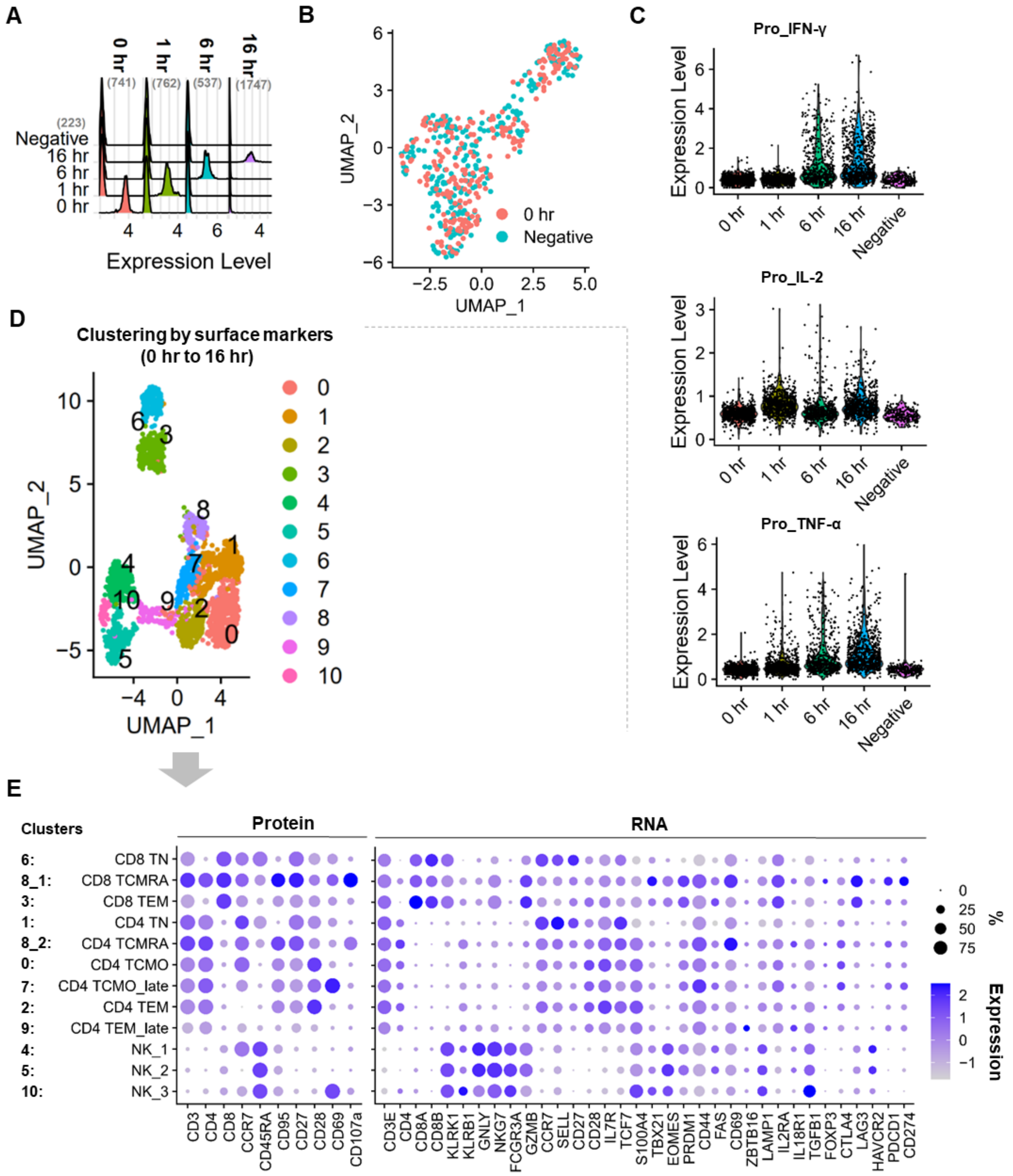


**Figure S4. Cells demultiplexing, clustering, and annotation**

(A) Cell samples demultiplexing based on the staining of anti-β2M-Oligo with the number of singlets bracketed. “Negative” cell group indicates cell sample without CapAbs staining. (B) Unsupervised clustering of cells with (0 hr) or without CapAbs staining (Negative). (C) Activation-dependent upregulation/downregulation of cytokine secretion. (D) Cell clustering based on the expression of a panel of surface protein markers as listed in (E). (E) Manual cell type annotation based on the expression profile of surface protein markers and a panel of genes relating to immune cell activation/differentiation.

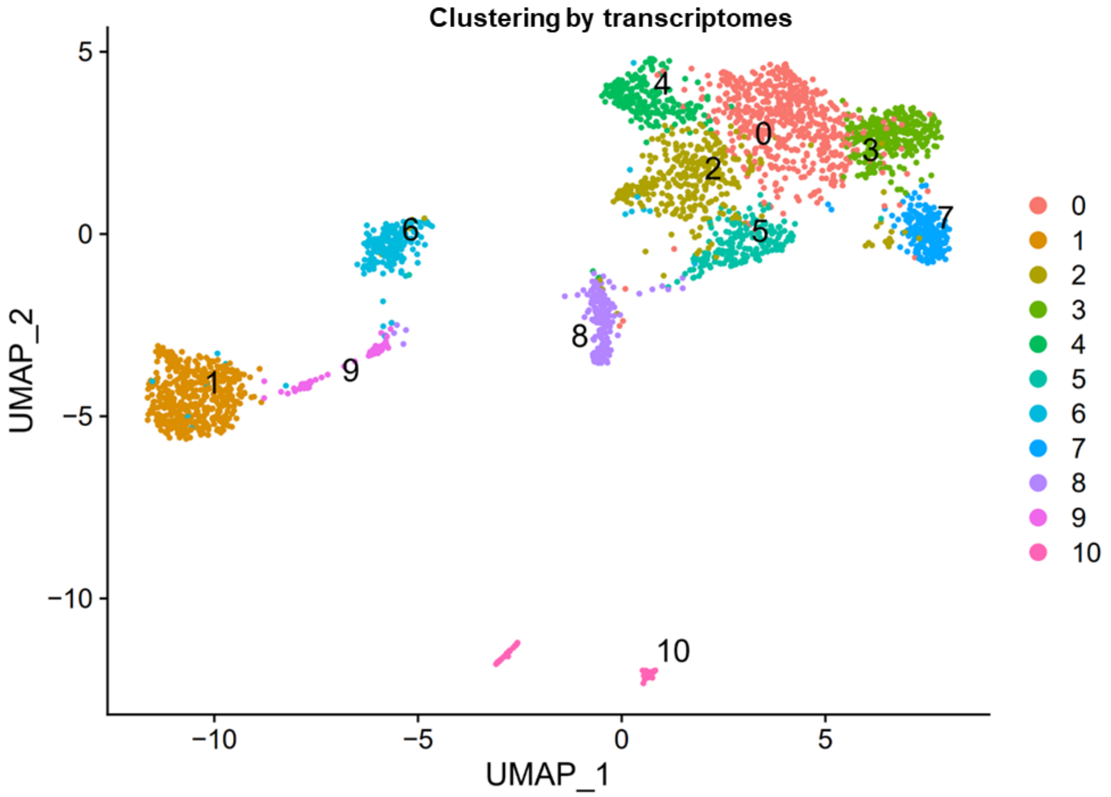
Clustering by transcriptomes
UMAP_2
UMAP_1
5
0
−5
−10
−10
−5
0
5
0
1
2
3
4
5
6
7
8
9
10

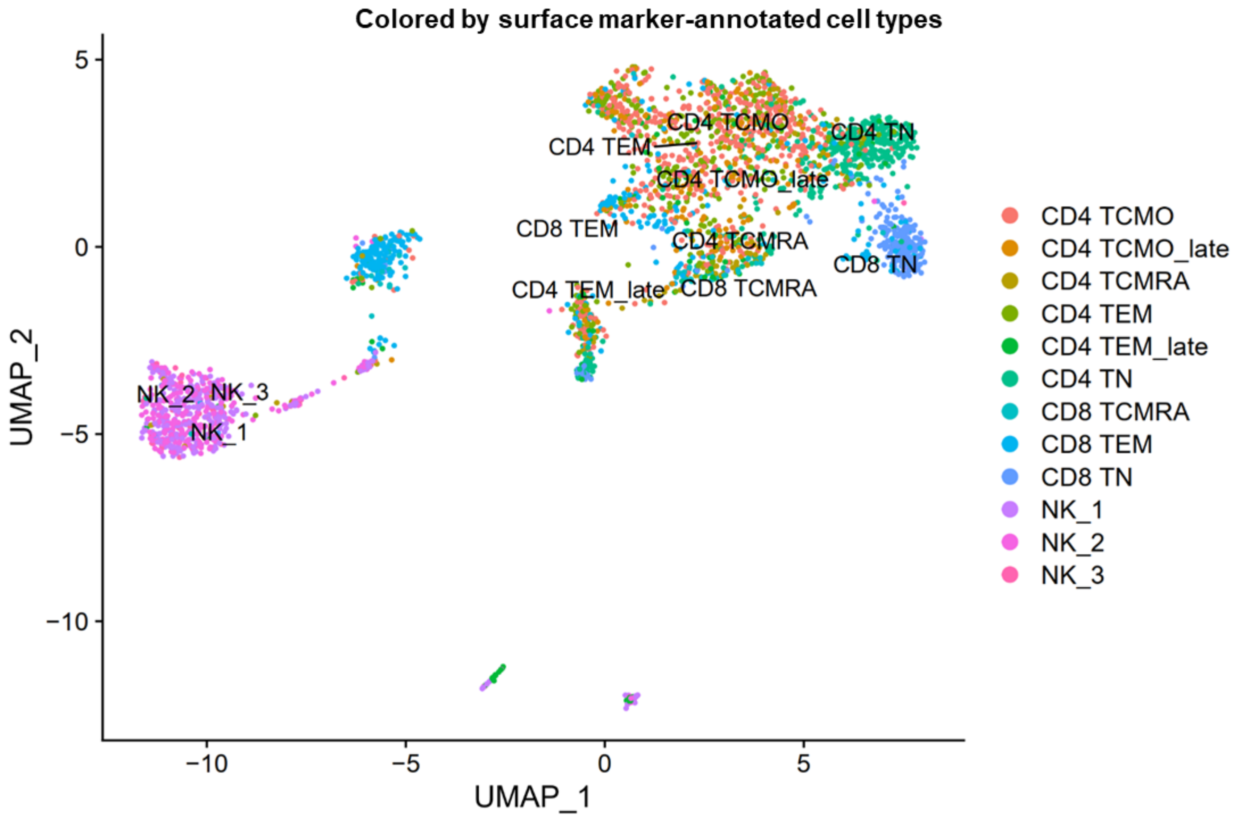
Colored by surface marker-annotated cell types
UMAP_2
UMAP_1
5
0
−5
−10
−10
−5
0
5
CD4 TCMO
CD4 TEM
CD4 TN
CD4 TCMO_late
CD8 TEM
CD4 TCMRA
CD8 TN
CD4 TEM_late
CD8 TCMRA
NK_2
NK_3
NK_1
CD4 TCMO
CD4 TCMO_late
CD4 TCMRA
CD4 TEM
CD4 TEM_late
CD4 TN
CD8 TCMRA
CD8 TEM
CD8 TN
NK_1
NK_2
NK_3

**Figure S5. Transcriptome profile-based cell clustering**

Cells were clustered by their gene expression profiles (top) and colored by cell subtypes (bottom) which were manually annotated based on phenotypic protein expression and a panel of selected genes.

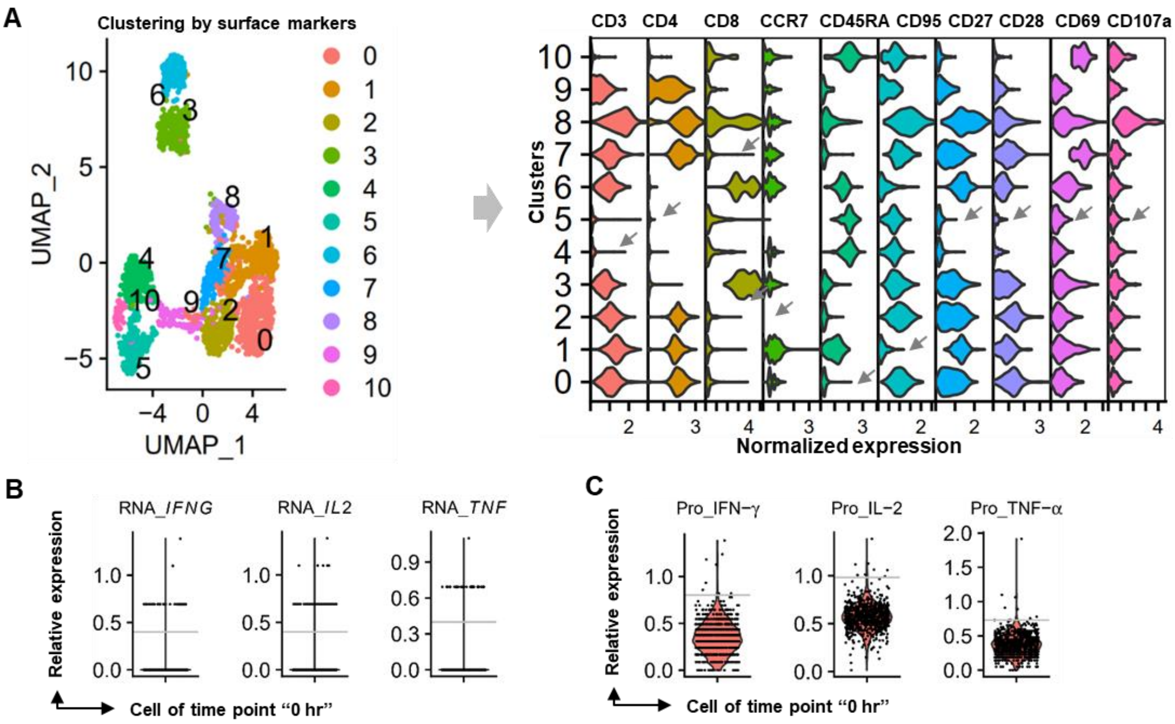


**Figure S6. Background determination of mRNA and protein expression**

(A-C) For the background staining of surface markers, cells were first analyzed by identifying clusters based on their phenotypic protein expression (the same to Figure S4D), and cell cluster (indicated by arrow) expected to have baseline expression of that marker was used for background subtraction, correspondingly (A). Cytokine gene expression (also surface-marker gene expression) threshold was set to clearly separate positive and negative cell groups (B) and cytokine secretion background was determined according to 99 quantiles of their baseline staining (before activation) (C).

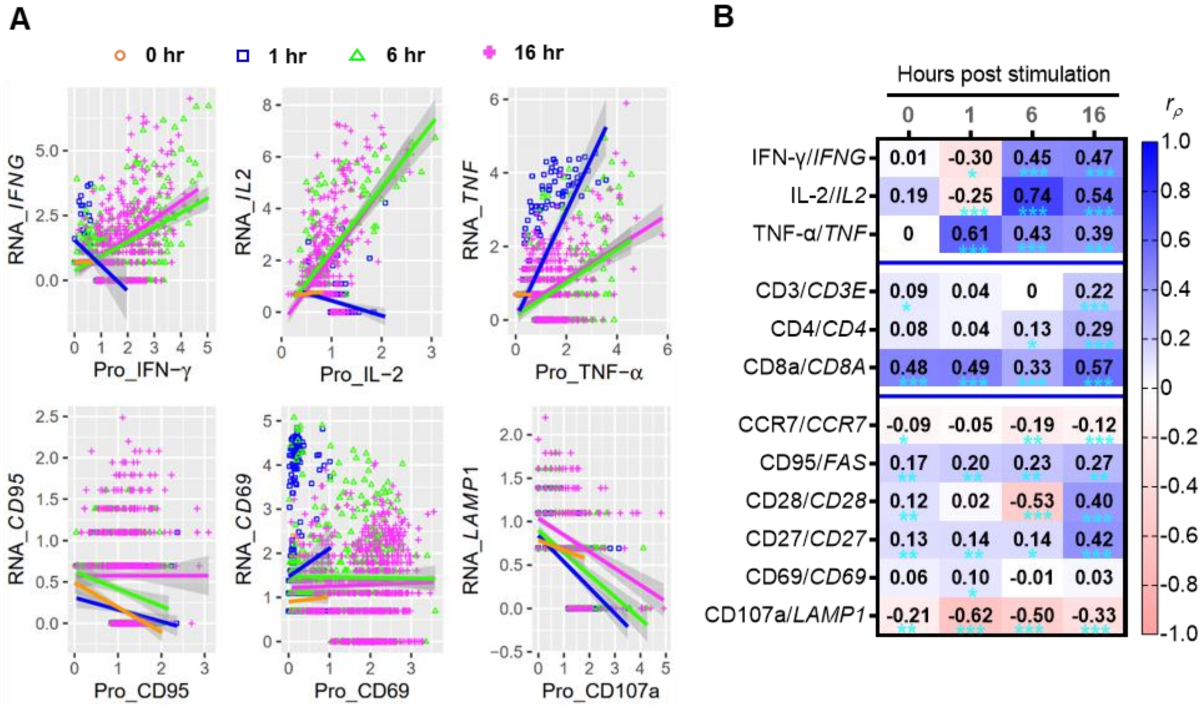


**Figure S7. Correlation analysis of protein expression with transcript abundance**

(A) Shown scattering plots were representative of the relationship of cytokine secretion versus cytokine gene level (top panel) and cell surface-born activation marker versus their gene expression (bottom panel). (B) A summary of Pearson correlation for paired protein/gene expression. *$P < .05$, **$P < .01$, ***$P < .001$.

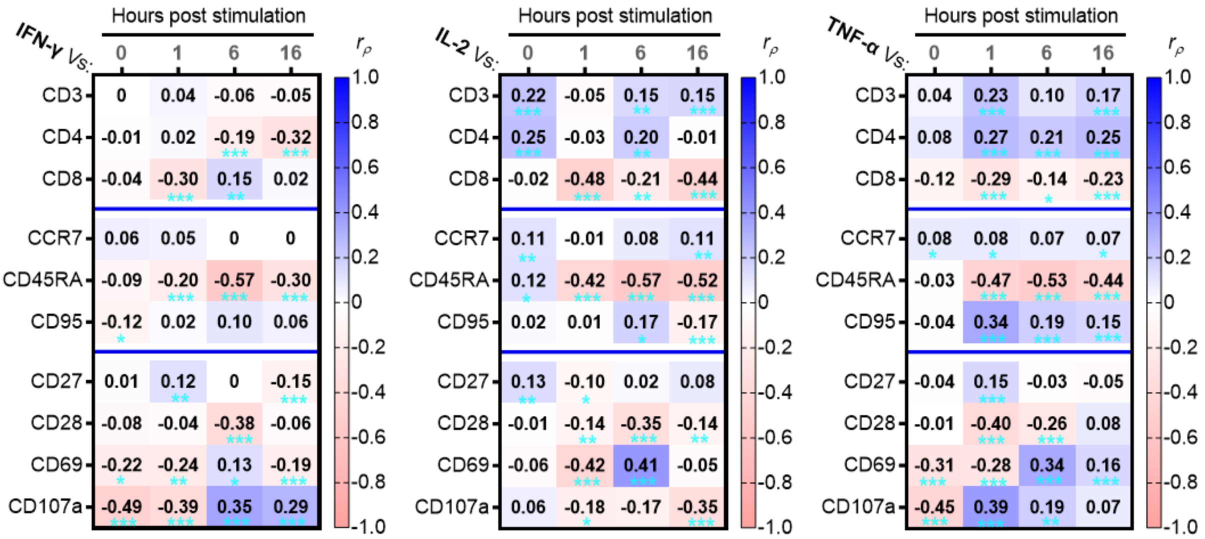


**Figure S8. Correlation analysis of cytokine secretion with phenotype/activation state**

Pearson correlation of cytokine release and phenotypic marker expression. *$P < .05$, **$P < .01$, ***$P < .001$.

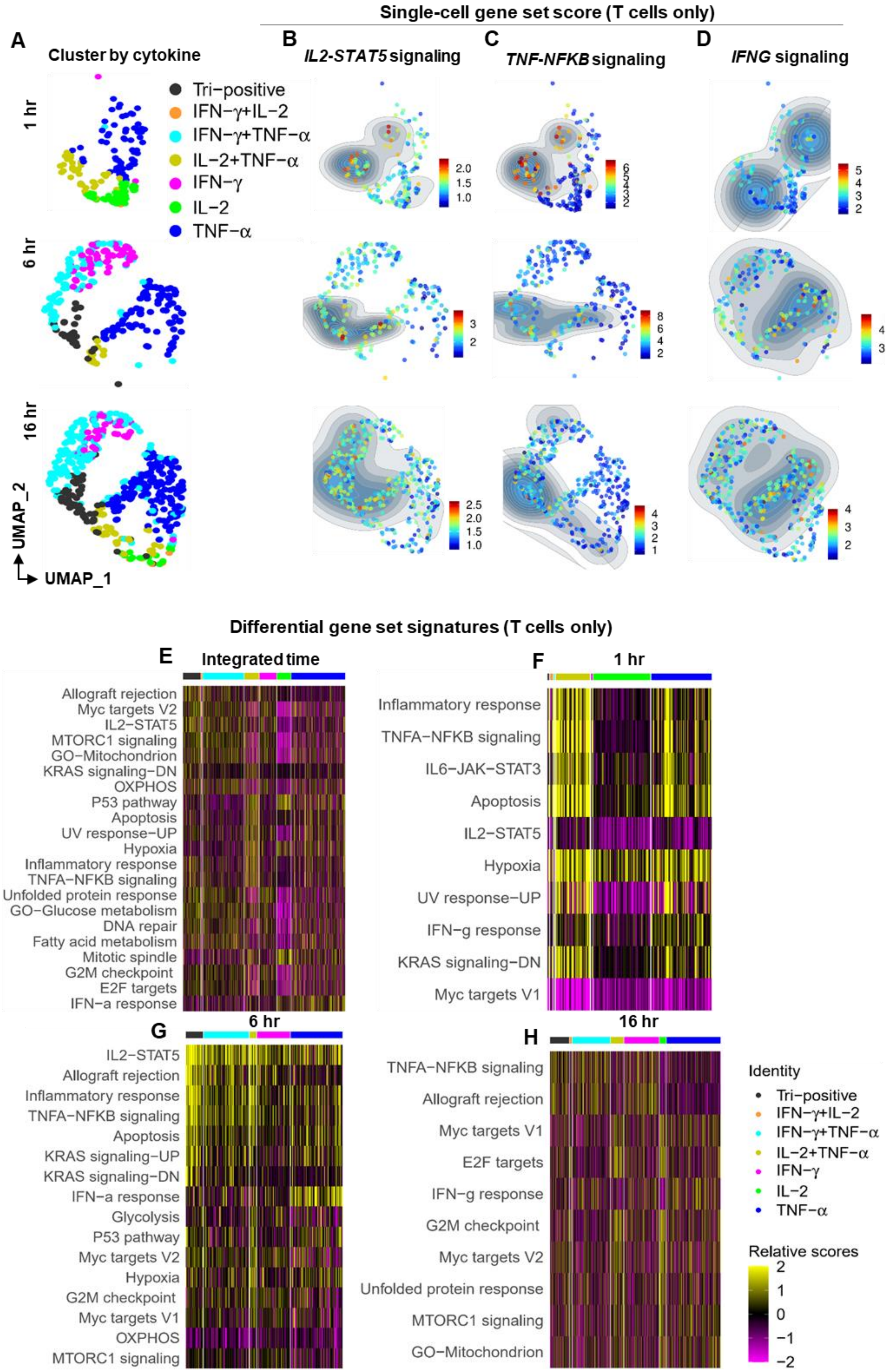

Single-cell gene set score (T cells only)
A
Cluster by cytokine
B
IL2-STAT5 signaling
C
TNF-NFKB signaling
D
IFNG signaling
1 hr
6 hr
16 hr
Tri−positive
IFN−γ+IL−2
IFN−γ+TNF−α
IL−2+TNF−α
IFN−γ
IL−2
TNF−α
UMAP_2
UMAP_1
Differential gene set signatures (T cells only)
E
Integrated time
Allograft rejection
Myc targets V2
IL2−STAT5
MTORC1 signaling
GO−Mitochondrion
KRAS signaling−DN
OXPHOS
P53 pathway
Apoptosis
UV response−UP
Hypoxia
Inflammatory response
TNFA−NFKB signaling
Unfolded protein response
GO−Glucose metabolism
DNA repair
Fatty acid metabolism
Mitotic spindle
G2M checkpoint
E2F targets
IFN−a response
F
1 hr
Inflammatory response
TNFA−NFKB signaling
IL6−JAK−STAT3
Apoptosis
IL2−STAT5
Hypoxia
UV response−UP
IFN−g response
KRAS signaling−DN
Myc targets V1
G
6 hr
IL2−STAT5
Allograft rejection
Inflammatory response
TNFA−NFKB signaling
Apoptosis
KRAS signaling−UP
KRAS signaling−DN
IFN−a response
Glycolysis
P53 pathway
Myc targets V2
Hypoxia
G2M checkpoint
Myc targets V1
OXPHOS
MTORC1 signaling
H
16 hr
TNFA−NFKB signaling
Allograft rejection
Myc targets V1
E2F targets
IFN−g response
G2M checkpoint
Myc targets V2
Unfolded protein response
MTORC1 signaling
GO−Mitochondrion
Identity
Relative scores
2
1
0
−1
−2


245

**Figure S9. Characterization of secretion polyfunctionality-associated gene set signatures**

(A-D) T cells with positive cytokine secretion were clustered according to their secretion profiles by time (A). Shown also were representative single-cell signature scores directly associated with the cytokine signaling over time (B-D). (E-H) Differential analysis of gene set signatures distinguishing T cells of varied cytokine-secreting potency. Shown were the top 7 gene set signatures with significant differences over cell groups compared.

**Table S1. List of antibodies used in this study**

| Antibody | Clone | Company | Catalog number |
|---|---|---|---|
| IFN-γ CapAb | *With the kit | Miltenyi Biotec | 130-090-433 |
| IL-2 CapAb | *With the kit | Miltenyi Biotec | 130-090-487 |
| TNF-α CapAb | *With the kit | Miltenyi Biotec | 130-091-267 |
| FITC-anti-IFN-γ | *With the kit | Miltenyi Biotec | 130-090-433 |
| PE-anti-IL-2 | *With the kit | Miltenyi Biotec | 130-090-487 |
| APC-anti-TNF-α | *With the kit | Miltenyi Biotec | 130-091-267 |
| FITC-msIgG1 | IS5-21F5 | Miltenyi Biotec | 130-113-761 |
| PE-msIgG2a | S43.10 | Miltenyi Biotec | 130-113-834 |
| APC-msIgG1 | IS5-21F5 | Miltenyi Biotec | 130-113-758 |
| BUV395-anti-CD3 | UCHT1 | BD Biosciences | 563546 |
| BV786-anti-CD4 | SK3 | BD Biosciences | 563877 |
| PE-Cy7-anti-CD8 | SK1 | BD Biosciences | 335787 |
| FITC-anti-CD19 | V CD19.11 | BD Biosciences | 555412 |
| PE-anti-CD45 | HI30 | BD Biosciences | 555483 |
| Purified anti-CD45 | HI30 | BioLegend | 304002 |
| TotalSeq-A0391 anti-CD45 | HI30 | BioLegend | 304064 |
| TotalSeq-A0251 anti-β2M | LNH-94 | BioLegend | 394601 |
| TotalSeq-A0252 anti-β2M | LNH-94 | BioLegend | 394603 |
| TotalSeq-A0253 anti-β2M | LNH-94 | BioLegend | 394605 |
| TotalSeq-A0254 anti-β2M | LNH-94 | BioLegend | 394607 |
| TotalSeq-A0034 anti-CD3 | UCHT1 | BioLegend | 300475 |
| TotalSeq-A0045 anti-CD4 | SK3 | BioLegend | 344649 |
| TotalSeq-A0046 anti-CD8 | SK1 | BioLegend | 344751 |
| TotalSeq-A0148 anti-CCR7 | G043H7 | BioLegend | 353247 |
| TotalSeq-A0063 anti-CD45RA | HI100 | BioLegend | 304157 |
| TotalSeq-A0386 anti-CD28 | CD28.2 | BioLegend | 302955 |

**Table S1. List of antibodies used in this study (continued)**

| Antibody | Clone | Company | Catalog number |
|---|---|---|---|
| TotalSeq-A0154 anti-CD27 | O323 | BioLegend | 302847 |
| TotalSeq-A0156 anti-CD95 | DX2 | BioLegend | 305649 |
| TotalSeq-A0155 anti-CD107a | H4A3 | BioLegend | 328647 |
| TotalSeq-A0146 anti-CD69 | FN50 | BioLegend | 310947 |

**Table S2. List of custom oligo used for antibody modification, q-PCR, protein library preparation and sequencing primers**

| DNA oligo | Sequence | Modification |
|---|---|---|
| Anti-IFN-γ_barcode | CCTTGGCACCCGAGAATTCCAACGAGTAGCTCGAGACAAAAAAAAAAAAAAAAAAAAAAAAA*A*A | 5'-Biotin-TEG, 3'-phosphorothioation |
| Anti-IL-2_ barcode | CCTTGGCACCCGAGAATTCCACAATCCCTTCTAGACGAAAAAAAAAAAAAAAAAAAAAAAAA*A*A | 5'-Biotin-TEG, 3'-phosphorothioation |
| Anti-TNF-α_ barcode | CCTTGGCACCCGAGAATTCCAATTAACAGGGATCCCGAAAAAAAAAAAAAAAAAAAAAAAAA*A*A | 5'-Biotin-TEG, 3'-phosphorothioation |
| FAM-polyT | CTACACGACGCTCTTTTTTTTTTTTTTTTTTTTTTTTTTTTTT | 5'-FAM |
| Q-PCR_ forward primer | AATGATACGGCGACCACCGAGATCTACACTCTTTCCCTACACGACGCTC | |
| Q-PCR_ reverse primer | CAAGCAGAAGACGGCATACGAGATATGCTGTCCCTTGGCACCCGAGAATTCCA | |
| Surface protein_1st_ additive primer | CCTTGGCACCCGAGAATT*C*C | 3'-phosphorothioation |
| Hashtag_1st_ additive primer | GTGACTGGAGTTCAGACGTGTGC*T*C | 3'-phosphorothioation |
| Surface protein_2nd_ amplification primer | CAAGCAGAAGACGGCATACGAGATCGTGATGTGACTGGAGTTCCTTGGCACCCGAGAATTC*C*A | 3'-phosphorothioation |
| Hashtag_2nd_ amplification primer | CAAGCAGAAGACGGCATACGAGATCGAGTAATGTGACTGGAGTTCAGACGTGT*G*C | 3'-phosphorothioation |
